# Multi-omic landscape of Mn(Ⅱ) oxidation in *Achromobacter pulmonis* ss21: From multicopper oxidase to metabolic support electron transfer

**DOI:** 10.64898/2026.03.02.709024

**Authors:** Jie Yu, Zhijuan Liu, Hongjie Wang

## Abstract

Microbially mediated Mn(Ⅱ) oxidation plays a critical role in regulating the global Mn(Ⅱ) cycle and represents an environmentally friendly strategy for remediation Mn(Ⅱ) contaminated waters. This study presents the first demonstration that *Achromobacter pulmonis* ss21, a bacterium isolated from Baiyangdian Lake, exhibits the excellent capacity to oxidze Mn(Ⅱ). The Mn(Ⅱ) oxidation efficiency of ss21 reached 98.82% and 97.05% for 200 and 400 mg/L Mn(Ⅱ), respectively. Transcriptome analysis revealed that direct Mn(Ⅱ) oxidation was catalyzed by genes encoding copper resistance system multicopper oxidase (*HV701_RS04390*), LLM-type flavin oxidoreductase (*HV701_RS19365*) and quinone oxidoreductase (*HV701_RS24690*), which regulate extracellular electron transfer for continuous Mn(Ⅱ) oxidation. In addition, thioredoxin (*HV701_RS19360*) and glutathione peroxidase (*HV701_RS19445*) genes maintained intracellular redox homeostasis, ensuring stable and efficient direct Mn(Ⅱ) oxidation under high Mn(Ⅱ) stress. Moreover, genes (*iscU*, *hscA*, *fliS*, *HV701_RS03300*, and *HV701_RS06395*) associated with metabolic support, motility, and transcriptional regulation supported indirect Mn(Ⅱ) oxidation. Metabolomics analysis revealed the upregulation of L-Tyrosine, L-Isoleucine, glutamic acid, Gln-His-His, Flavin Adenine Dinucleotide (FAD), xanthine related to ss21 Mn(Ⅱ) oxidation, which corresponded to the direct Mn(Ⅱ) oxidation genes. This study provides a comprehensive understanding of the molecular mechanisms of biological Mn(Ⅱ) oxidation by *Achromobacter sp*. and highlights its potential application in the bioremediation of Mn contaminated aquatic environments under high metal stress conditions.

**IMPORTANCE:** Microbially mediated Mn(Ⅱ) oxidation is a fundamental process in global biogeochemical cycles and offers a sustainable pathway for remediating heavy metal-polluted waters. *Achromobacter pulmonis* ss21 showed excellent performance in Mn(Ⅱ) oxidation. The highly efficient removal of Mn(Ⅱ) was achieved through oxidoreductase catalysis, regulation of extracellular electron transfer, maintenance of redox homeostasis, and ensurance of stable and efficient Mn(Ⅱ) oxidation under high Mn(Ⅱ) stress. Moreover, the Mn(Ⅱ) oxidation was supported by metabolites, which prevented irreversible protein damage from oxidative stress induced by high Mn(Ⅱ) concentration, alleviating oxidative stress, and stimulating the production of ROS. These findings expand the known diversity of Mn(Ⅱ) oxidizing bacteria and offer valuable information for the molecular mechanisms of biological Mn(Ⅱ) oxidation.

## INTRODUCTION

Manganese-oxidizing bacteria (MOB) are widely distributed across diverse environments, including marine, freshwater, and terrestrial ecosystems (1). MOB was characterized by the crucial effect on the Mn(Ⅱ) biogeochemical cycling, and significantly influence the fate of various contaminants in the ecosystem (2). The MOB exhibit different Mn(Ⅱ) oxidation capabilities at various Mn(Ⅱ) concentration. Certain bacteria had an excellent capacity for the lower Mn(Ⅱ) concentration conditions (< 55 mg/L). For example, the *Bacillus altitudinis* BYD19 could reduce Mn(Ⅱ) from 51.00 to 3.50 mg/L, and the *Cupriavidus* sp. HY129 achieved the highest Mn(Ⅱ) removal rate approximately 95.79% at the 5 mg/L initial Mn(Ⅱ) concentration (3, 4). In a higher Mn(Ⅱ) concentration condition (> 55 mg/L) the MOB extracted from Mn(Ⅱ)-rich niches have demonstrated the extraordinary Mn(Ⅱ) tolerance (5). The Mn(Ⅱ) oxidation activity of the *Lysinibacillus xylanilyticus* M125 reached the optimum value at Mn(Ⅱ) concentration of 660 mg/L (6). Moreover, the *Bacillus thuringiensis* H38 and the *Aeromonas hydrophila* DS02 achieved the highest Mn(Ⅱ) oxidation rate at the initial Mn(Ⅱ) concentration of 550 mg/L (7, 8). The *B. zhangzhouensis* exhibited optimal growth and Mn(Ⅱ) oxidation efficiency at Mn(Ⅱ) concentration of 1000 mg/L (9). The *Bacillus altitudinis* HM-12 maintained a 90.6% Mn(Ⅱ) removal efficiency at an extreme Mn(Ⅱ) concentration of 5500 mg/L (10).

The predominant biological oxidation mechanism for Mn(Ⅱ) is directly mediated by multicopper oxidases (MCOs) and animal heme peroxidase (AHPs) (11, 12). MCOs catalyze the oxidation of Mn(Ⅱ) to Mn(Ⅲ), which is subsequently transformed into MnO_2_ (13, 14). Recent research has demonstrated the model strain utilized MCOs to oxidize Mn(Ⅱ), such as *Bacillus pumilus* WH4, *Pseudomonas putida* GB-1, *Leptothrix discophora* SS-1 (12, 15, 16). Several MCOs including MnxG, MofA, CotA and MoxA acted as the active Mn(Ⅱ) oxidases (16). The *Bacillus pumilus* WH4 oxidized Mn(Ⅱ) through the catalysis of *cotA* gene, and strain engineered to overexpress the CotA exhibited enhancement of Mn(Ⅱ) removal capability (16). Similarly, *Leptothrix discophora* SS-1 utilizes MofA and MnxG to drive Mn(Ⅱ) oxidation (17). In addition, it was found that disruption of MoxA through insertional mutagenesis will lead to the loss of Mn(Ⅱ) oxidation ability, indicating MoxA is the key enzyme catalyzing Mn(Ⅱ) oxidation (18). Bacteria that utilizd AHPs to oxidize Mn(Ⅱ) were isolated from various environmental conditions (19, 20). The *MopA* isolated from bacteria belongs to the AHP family, which was essential in Mn(Ⅱ) biological oxidation (21). Genetic and structural studies have confirmed that *MopA* in *Erythrobacter sp.* SD-21 and *Aurantimonas manganoxydans* SI85-9A1, promoting the oxidation of Mn(Ⅱ) to Mn(Ⅳ) through the Mn(Ⅲ) intermediate, and the *MopA* gene does not express in the absence of Mn(Ⅱ) (19, 21). Nevertheless, the recognition of microbial Mn(Ⅱ) oxidation mechanisms was insufficient that limited to directly synthesizing oxidase enzymes.

Microorganisms altered their surrounding environment through metabolic activities, triggering indirectand non-enzymatic Mn(Ⅱ) oxidation mechanism. By producting extracellular superoxide radicals (O_2_**^·^**^−^), the strain rapidly degraded citrate and efficiently mediated the indirectly oxidation Mn(Ⅱ) oxidation (22). Extracellular polymeric substances (EPS) efficiently activated bacterial cell generated O_2_^·-^, through electron transfer mediated by its internal NAD(P)H oxidase and non-enzymatic components, significantly enhanced Mn(Ⅱ) oxidation capacity (23). Moreover, the Mn(Ⅱ) oxidation process involved the cooperation of directly and indirectly genes. The MCOs and genes that coding copper homeostasis proteins, cytochrome c oxidase as the directly Mn(Ⅱ) oxidation genes, which was linked to the indirectly genes coding nitrogen cycling and electrons transfer, synergeticly drived Mn(Ⅱ) oxidation (24). Therefore, Mn(Ⅱ) oxidation is indeed a complex and multi-layered process, involving redox dynamics and environmental factors.

In this study, the Mn(Ⅱ) oxidizing bacterium *Achromobacter* ss21 was isolated from Baiyangdian sediments, which shown significant capability for Mn(Ⅱ) oxidation. The effects of initial Mn(Ⅱ) concentration, pH, and temperature on Mn(Ⅱ) oxidation by *Achromobacter* ss21 were studied, and the molecular mechanisms of Mn(Ⅱ) oxidation were proposed by transcriptome and metabolome analyses. This study will provide deeper insight into Mn(Ⅱ) oxidation by bacteria, and be benefit to understand the global Mn(Ⅱ) cycle.

## MATERIALS AND METHODS

### STRAIN ISOLATION AND GENOME SEQUENCING

*Achromobacter pulmonis* ss21 was procured from sediment samples collected from Baiyangdian Lake, Xiong’an New Area, Hebei Province. The strain ss21 was incubated in the dark using the sterile Luria-Bertani (LB) medium (5.0 g/L peptone, 5.0 g/L NaCl, and 2.5 g/L yeast extract) at 30°C with 170 rpm shaking. To qualitate the Mn(Ⅱ) oxidation function of ss21 strain, the cultured solution was mixed with Leucoberbelin blue (LBB) reagent in a 1:1 ratio to observe the color change of solution color. Genomic sequencing of strain ss21 was performed based on 16S rRNA gene sequencing. The 16S rRNA gene was amplified by polymerase chain reaction (PCR) using universal bacterial primers 27F (AGAGTTTGATCCTGGCTCAG) and 1492R (GGTTACCTTGTTACGACTT). The resulting PCR products were subjected to paired-end sequencing (2 × 150 bp) on the Illumina sequencing platform. The obtained sequences were compared with reference sequences in the GenBank RefSeq database using BLAST for taxonomic identification. Phylogenetic trees of ss21 were constructed using MEGA 7.0 software.

### MN(Ⅱ) OXIDATION ASSAYS

To verify the ability of ss21 strain to oxide Mn(Ⅱ), ss21 was inoculated at 1% (v/v) inoculum into erlenmeyer flasks containing 100 mL LB medium, and the initial Mn(Ⅱ) concentrations were set at 50, 100, 150, 200, 400, 600, and 800 mg/L, respectively. These erlenmeyer flasks were cultured at 30°C with 170 rpm shaking. To investigate the influence of temperatureon Mn(Ⅱ) oxidation, the inoculation temperature was set at 10, 20, 25, 30, and 35 °C, respectively. To evaluate the effects of initial pH on Mn(Ⅱ) oxidation, the pH of LB media was adjusted to 5, 6, 7, and 8, respectively, the begin of ss21 strain inoculated with Mn(Ⅱ) during the incubation period, samples were collected daily to determine residual Mn(Ⅱ) concentrations and density at 600 nm (OD_600_). To explore the adsorption and oxidation fraction of Mn(Ⅱ) by ss21, a four-step sequential extraction procedure was employed (Tang et al., 2016). The Mn concentration was divided into four states: residue, extracellularly adsorption, intracellularly adsorption, and oxidation. The detailed extraction procedures of each state are provided in Text S1. All experiments were conducted in triplicate to ensure reproducibility. Besides the temperature influence experiment, all erlenmeyer flasks were cultured at 30°C with 170 rpm shaking.

### TRANSCRIPTOME AND METABOLOMES ANALYSES

The ss21 strain was cultured in 200 mL of LB medium with different initial Mn(Ⅱ) concentrations, the cells cultured without Mn(Ⅱ) were as the control group, bacterial samples were collected at 10 h and 48 h for metabolomic and transcriptomic analyses, collected at 10 and 48 h. For transcriptome analysis, the total RNA was extracted form the collected strain samples using the MJZol total RNA extraction kit, and the total RNA libraries were constructed by the Illumina Stranded Total RNA Prep Kit. The quantification of gene was performed by sequencing of libraries. The GO and KEGG enrichment analysis were used to functional annotation of Mn(Ⅱ) oxidation. For metabolomes analysis, the metabolites of ss21 were identified using liquid chromatography-mass spectroscopy (LC-MS) with UHPLC-Q Exactive HF-X (ThermoFisher Scientific, USA). The raw data from LC-MS was edited by Progenesis QI (Waters Corporation, Milford, USA) software, then, the edited data were imported to Majorbio database to identifiy the metabolites of ss21. The PCA, KEGG pathway, KEGG topology and clustered heatmaps analyses were conducted to identify the function of different metabolites.

### ANALYTICAL METHODS

The morphology and elemental composition analyses were performed using scanning electron microscopy and energy dispersive spectroscopy (SEM-EDS, ZEISS Sigma 360, Germany). The crystalline phases of Mn in biogenic manganese oxides (BioMnOx) were detected using X-ray diffraction analysis (XRD, Rigaku SmartLab SE, Japan). The valence states of Mn were analyzed using X-ray photoelectron spectroscopy (XPS, Thermo Scientific K-Alpha, USA), and the BioMnOx samples was harvested at 10, 48, 72, and 96 h for XPS. The concentration of Mn(Ⅱ) was determined using Atomic Absorption Spectrometry (AAS, SKYRAY AAS900, China). The pH was measured using desktop acidity meter. The OD_600_ of ss21 was measured using a UV-Vis spectrophotometer (Hitachi U-3010, Japan).

## RESULTS AND DISCUSSION

### IDENTIFICATION OF STRAIN

Strain ss21 were isolated and purified from Baiyangdian Lake sediments, and subsequently screened for Mn(Ⅱ) oxidizing capabilities. The formation of brown precipitate was observed after 6 d when the strain inoculated with Mn(Ⅱ) (Fig. S1a). Then, the ss21 strain were inoculated into liquid LBB medium containing 40 mg/L Mn(Ⅱ), the bright blue color of LBB medium confirmed the presence of BioMnOx, whereas, no color change was observed in the control group without Mn(Ⅱ) supplementation (Fig. S1b). These results indicated that the production of BioMnOx by ss21 strain. Based on 16S rRNA gene sequence, the complete genome of ss21 was determined to be 6,285,168 bp in length, comprising one circular chromosome (Fig.1a). Phylogenetic analysis indicated that this strain shared high similarity with the sequence date of *Achromobacter pulmonis*, with a 16S rRNA sequence similarity of 99.70% (Fig.1b). Further genomic comparative analysis shown ss21 has the 99.69% homology with *Achromobacter pulmonis* R-16442 (Fig.1c). Consequently, this Mn(Ⅱ) oxidizing strain was designated as *Achromobacter pulmonis* ss21.

**Fig. 1.**
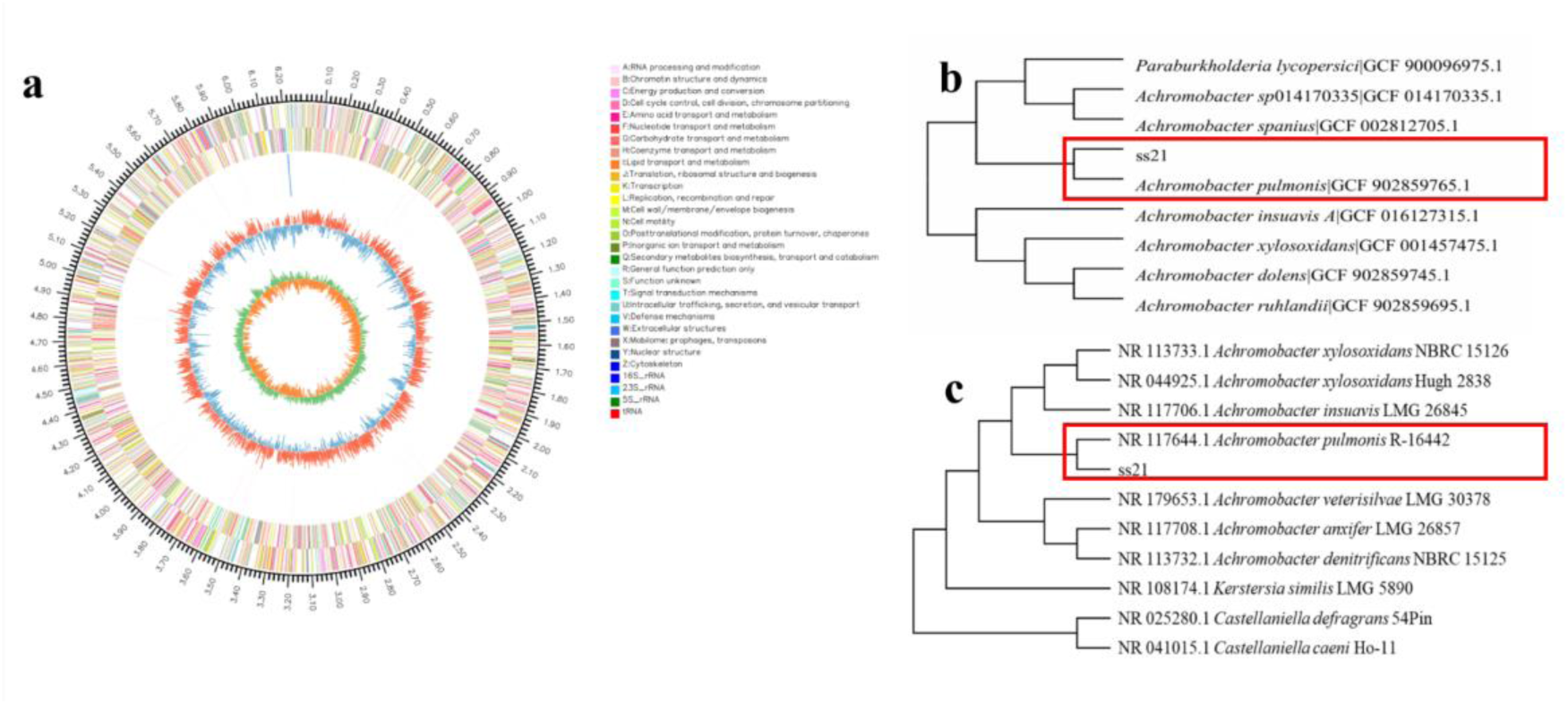
Genome assembly and identification of strain ss21(a), Phylogenetic analysis of ss21 (b and c).

### DETERMINATION OF MN(Ⅱ) OXIDATION CAPACITY OF STRAIN

The Mn(Ⅱ) oxidation capacity of bacterial ss21 under different factors were systematically analyzed (Fig. 2). The Mn(Ⅱ) removal efficiency reached 83.40%, 84.39%, 91.93%, 98.82%, 97.05%, 42.55%, and 54.73%, when initial Mn(Ⅱ) concentration was 50, 100, 150, 200, 400, 600, and 800 mg/L, respectively (Fig. 2a). Further more, Mn(Ⅱ) oxidation showed an initial increase followed by a decrease with the rising Mn(Ⅱ) concentration. At 50 mg/L Mn(Ⅱ) concentration, the Mn(Ⅱ) oxidation efficiency is relatively low, at 83.40%. This may be attributed to the lower Mn(Ⅱ) concentration activated the bacterial oxidation capability insufficient (25). The highest Mn(Ⅱ) oxidation efficiency of 98.82% was observed at the 200 mg/L initial Mn(Ⅱ), where the Mn(Ⅱ) concentration decreased from 224 to 2.65 mg/L. However, when the Mn(Ⅱ) concentration exceeded 600 mg/L, the Mn(Ⅱ) oxidation efficiency decreased significantly. Excessive Mn(Ⅱ) obviously inhibited the bacterial growth and Mn(Ⅱ) oxidation rate (6). The ss21 strain growth entered a rapid exponential phase before 24 h cultivation, and the OD_600_ reached approximately 1.34 at 24 h across all initial Mn(Ⅱ) concentrations (Fig. S2a). This results revealed that the ss21 strain had the excellent capability of high Mn(Ⅱ) concentration oxidation and resistance. The pH of the medium gradually increased from 5.78 to 9.0, and stabilizing at alkaline conditions at 48 h (Fig. S2b). However, the final pH values varied under different initial Mn(Ⅱ) concentrations. At relatively low Mn(Ⅱ) concentration (50 - 200 mg/L), the pH reached approximately 8.8 - 9.1, while at higher Mn(Ⅱ) concentration (> 200 mg/L), the pH stabilized around 8.0 - 8.5. These results confirmed that alkaline conditions are optimal for the growth and Mn(Ⅱ) oxidation of strain ss21. At around 200 mg/L Mn(Ⅱ), the metabolic activity and pH adjusting ability of strain were maximized, resulting in the highest Mn(Ⅱ) oxidation efficiency.

**Fig. 2.**
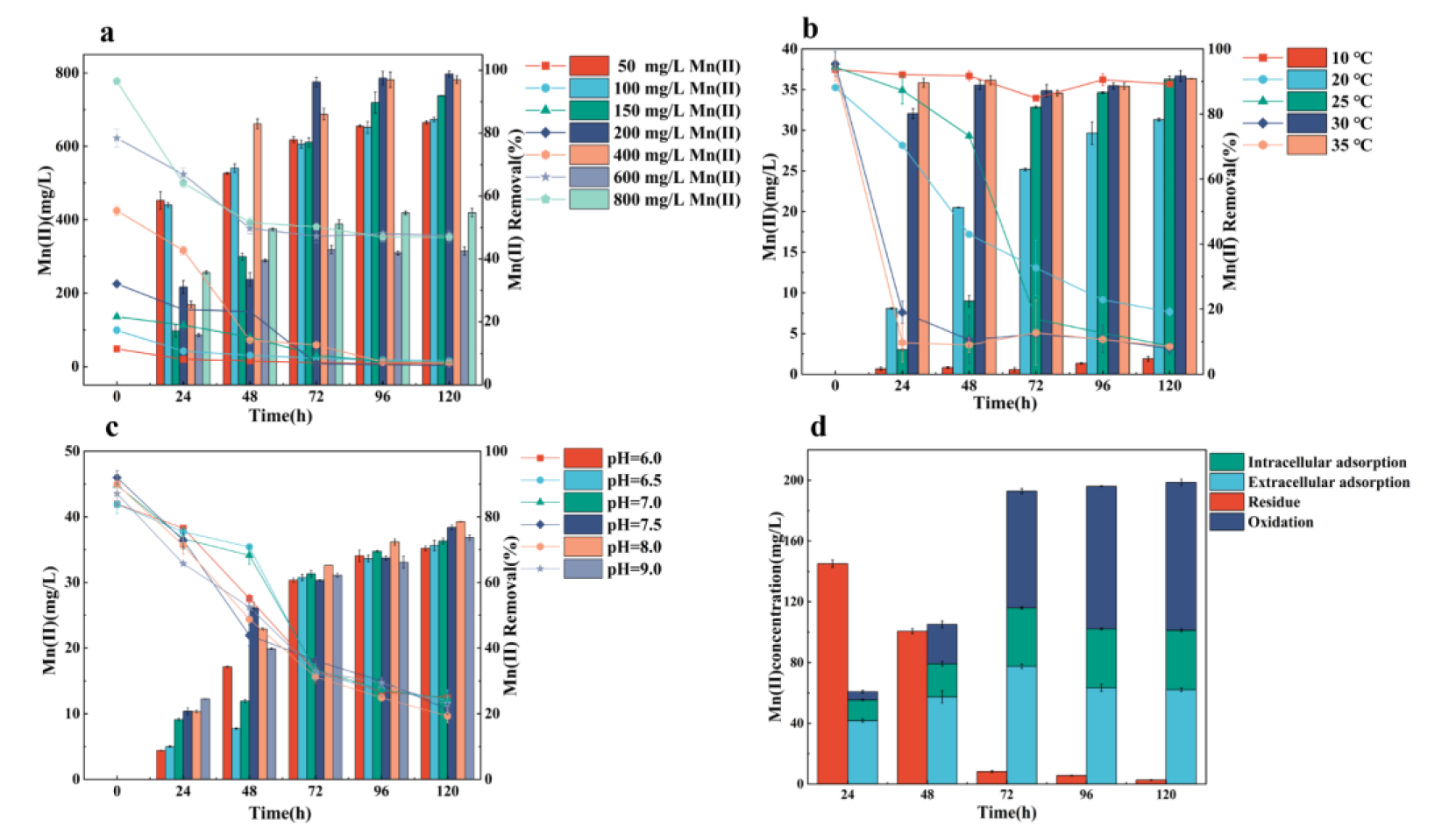
Change in Mn(Ⅱ) concentration under different initial Mn(Ⅱ) concentration (a), temperature (b), pH (c), Mn(Ⅱ) adsorption and oxidation by ss21 with initial Mn(Ⅱ) concentration of 200 mg/L (d).

The strain ss21 exhibited an increasing Mn(Ⅱ) oxidation efficiency under the different temperature (Fig. 1b). After 120 h, the Mn(Ⅱ) removal efficiency reached 4.78%, 78.27%, 90.70%, 91.69% and 90.87% at 10, 20, 25, 30 and 35℃, respectively. The ss21 showed the highest Mn(Ⅱ) removal efficiency at 30℃, with Mn(Ⅱ) concentration decreasing from 38.16 to 3.42 mg/L, and a higher OD_600_ of 1.56 was observed after 24 h of incubation at 30℃ (Fig. S2c). The pH increased to about 7.5 at 10°C, while increased to about 9.5 at higher temperatures (Fig. S2d). The low temperature suppressed the growth and metabolic activity rates of strain, while high temperature promoted the the critical enzymes and proteins expression of ss21, ultimately adjusting the pH of the medium to alkaline condition for the efficient Mn(Ⅱ) oxidation (8). The Mn(Ⅱ) removal efficiency of strain reached 70.34%, 71.29%, 72.62%, 76.76%, 78.51% and 73.66% at pH 6.0, 6.5, 7.0, 7.5, 8.0, and 9.0, respectively (Fig. 2c). The strain exhibited higher Mn(Ⅱ) removal performance and OD_600_ at pH 8, indicating that alkaline environment was more suitable for Mn(Ⅱ) oxidation process of ss21(Fig. S2e). The pH affected the activity of enzymes, thereby influencing bacterial growth and the oxidation rate of Mn(Ⅱ) (26), and the strong acid and alkali environment will decrease the efficiency of BioMnOx generation (4, 8).

Furthermore, Mn(Ⅱ) oxidation and adsorption activities by ss21 took different parts (Fig.2d). Before 48 h of cultivation, extracellular and intracellular adsorption major contribute to Mn(Ⅱ) removal, while the oxidation value of Mn(Ⅱ) was 25.83 mg/L at 48 h. Thereafter, the oxidation of Mn(Ⅱ) obvious increased to 76.94 mg/L at 72 h. These suggested that strain ss21 carried rapid growth within 24 h of cultivation, then carried efficient Mn(Ⅱ) oxidation from 48 h to 72 h, which were in accord with the result that OD_600_ reached maximum at 24 h (Fig. S2a). As the Mn(Ⅱ) oxidation activity gradually increased, extracellular and intracellular Mn(Ⅱ) adsorption decreased slightly from 72 h to 120 h. Therefore, it can be speculated that the strain ss21 adsorbed Mn(Ⅱ) from solution to outside and inside the cell membrane firstly, then continue came to transform the adsorbed Mn(Ⅱ) into BioMnOx.

### CHARACTERIZATION OF BIOMNOX

The morphology of ss21 strain was rod-shaped and have a rough surface, cover with noticeable granular and flocculent substances (Fig. 3a). Subsequently, the aggregated and clumped substances was observed on the the surface of the bacterial cells, which proved that the formation of BioMnOx by ss21 (Fig. 3b). BioMnOx were characterized by EDS, and it was found that Mn, O, C, and N elements in the mixed phase structure of BioMnOx and bacterial cells (Fig. 3c, d and e). The XRD characterization of BioMnOx showed characteristic peaks at 2θ values of 21.20°, 31.78°, 34.66°, 45.52°, and 56.58°, which indicated the coexistence of Mn(Ⅱ), Mn(Ⅲ), and Mn(Ⅳ) valence states, the BioMnOx were identified as a complex mixture primarily consisting of MnCO_3_, Mn_2_O_3_, and MnO_2_ (Fig. 3f). These results consistent with the above conclusion that Mn(Ⅱ) oxidation process of ss21 involved the adsorption of soluble Mn(Ⅱ), subsequent the strain utilized soluble Mn(Ⅱ) as electron acceptor and oxidated Mn(Ⅱ) into BioMnOx (27, 28). The XPS characterization of BioMnOx at different cultivation time revealed the consistent presence of Mn(Ⅱ), Mn(Ⅲ), and Mn(Ⅳ), with Mn(Ⅲ) was the predominant Mn species (Fig. 3g). The XPS spectra of the Mn 2p region exhibited two characteristic spin-orbit doublets, corresponding to the Mn 2p_3/2_ peak located at 641.78 eV and the Mn 2p_1/2_ peak located at 653.48 eV, respectively. The Mn 2p_3/2_ spectra could be split into three peaks with binding energies at 641.7 eV, 641.6 eV and 642 eV, representing Mn(Ⅱ), Mn(Ⅲ), Mn(Ⅳ), respectively (29, 30). From 10 h to 72 h, the proportion of Mn(Ⅱ) gradually decreased from 56.2% to 36.62%, while Mn(Ⅲ) increased from 40.13% to 52.20%, and Mn(Ⅳ) exhibited noticeable growth from 3.67% to 11.18%, indicating a Mn(Ⅱ) continuous oxidation process to form the BioMnOx. This observation is consistent with the general understanding that Mn(Ⅱ) oxidation proceeds through Mn(Ⅲ) as a key intermediate, and the resulting BioMnOx structures are typically mixed-valence Mn(Ⅲ/Ⅳ) oxides (31, 32).

**Fig. 3.**
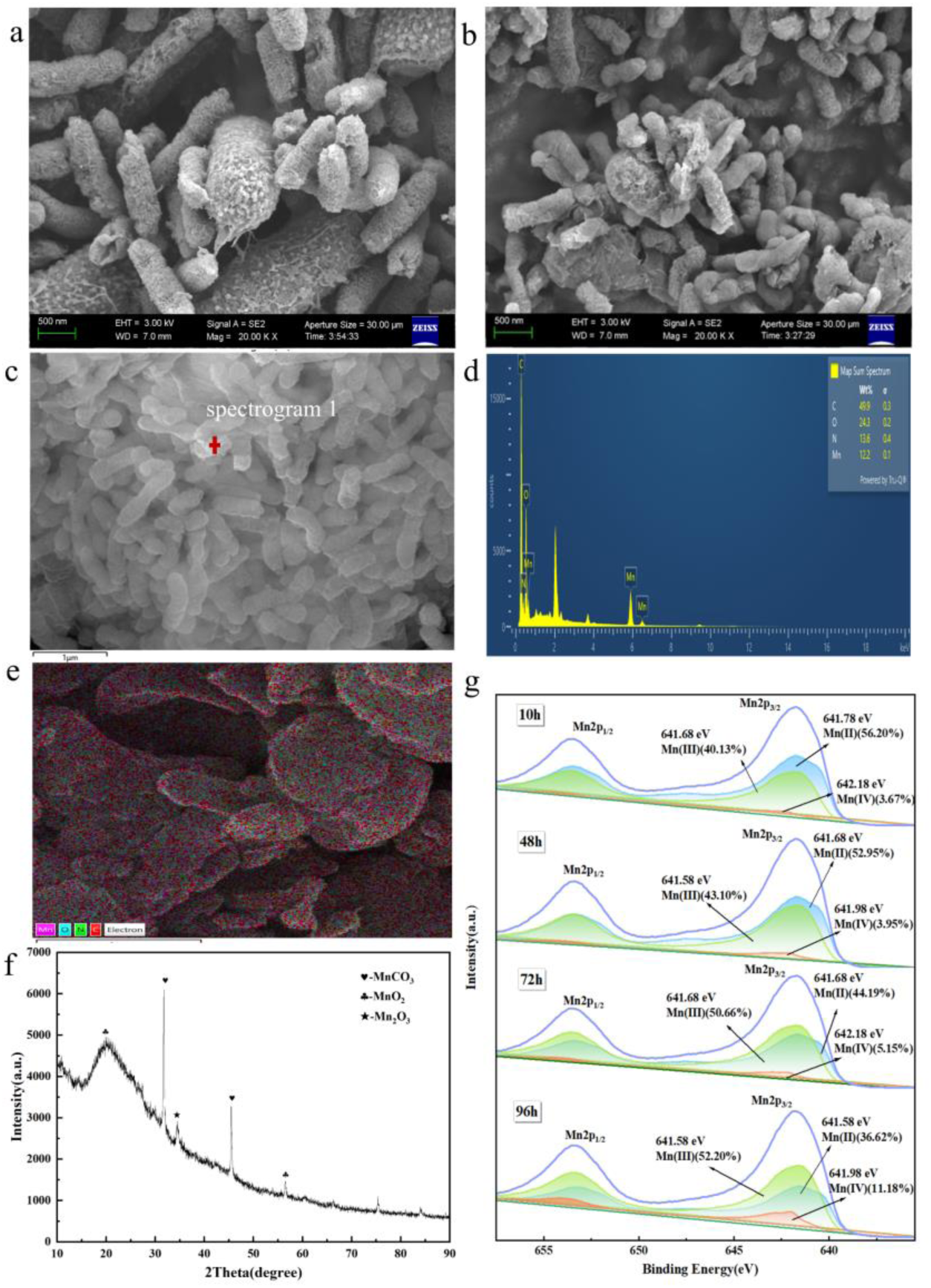
SEM image of ss21(a) and ss21 during oxidation process (b), EDS at region of spectrogram 1 (c and d), XRD patterns (f), XPS peak fitting of Mn 2p at different cultivation time (g).

### DIFFERENTIAL GENE EXPRESSION OF STRAIN

To further investigate the main functional genes of Mn(Ⅱ) oxidation process by ss21 strain, the transcriptomes were conducted in ss21 strain and ss21 with different Mn(Ⅱ) initial concentration (Fig. 4). In ss21 group, 583 genes were observed to be upregulated, while 1341 genes were downregulated, exhibiting 1924 differentially expressed genes, when compared 48 h with 10 h (Fig. 4a). At 10 h, the ss21exhibited 229, 308, and 611 differentially expressed genes in ss21 + 20 mg/L Mn(Ⅱ), ss21 + 60 mg/L Mn(Ⅱ) and ss21 + 200 mg/L Mn(Ⅱ) groups, respectively. These results showed that the differentially expressed genes decreased when ss21 strain cultivated with Mn(Ⅱ), and the differentially expressed genes increased with Mn(Ⅱ) initial concentration increased. At 48 h, the ss21 exhibited 903, 1055, and 2401 differentially expressed genes in ss21 + 20 mg/L Mn(Ⅱ), ss21 + 60 mg/L Mn(Ⅱ) and ss21 + 200 mg/L Mn(Ⅱ) groups, respectively. The number of differentially expressed genes were significantly increased at 48 h when compared to 10 h in the corresponding groups. Moreover, the differentially expressed genes increased with the rise of Mn(Ⅱ) concentrations. This scene was possibly due to the toxicity of Mn(Ⅱ) ions inhibited the gene expression of ss21, whereas the higher Mn(Ⅱ) initial concentration stimulated the gene expression of strain, which was beneficial for the oxidative ability for Mn(Ⅱ) by ss21. The gene expression of strain was intensely activated by in the presence of Mn(Ⅱ) from 10 h to 48 h, including these genes related to Mn(Ⅱ) converted to BioMnOx. Additionally, the ss21 strain possessed the excellent Mn(Ⅱ) removal capacity under higher Mn(Ⅱ) concentration.

**Fig. 4.**
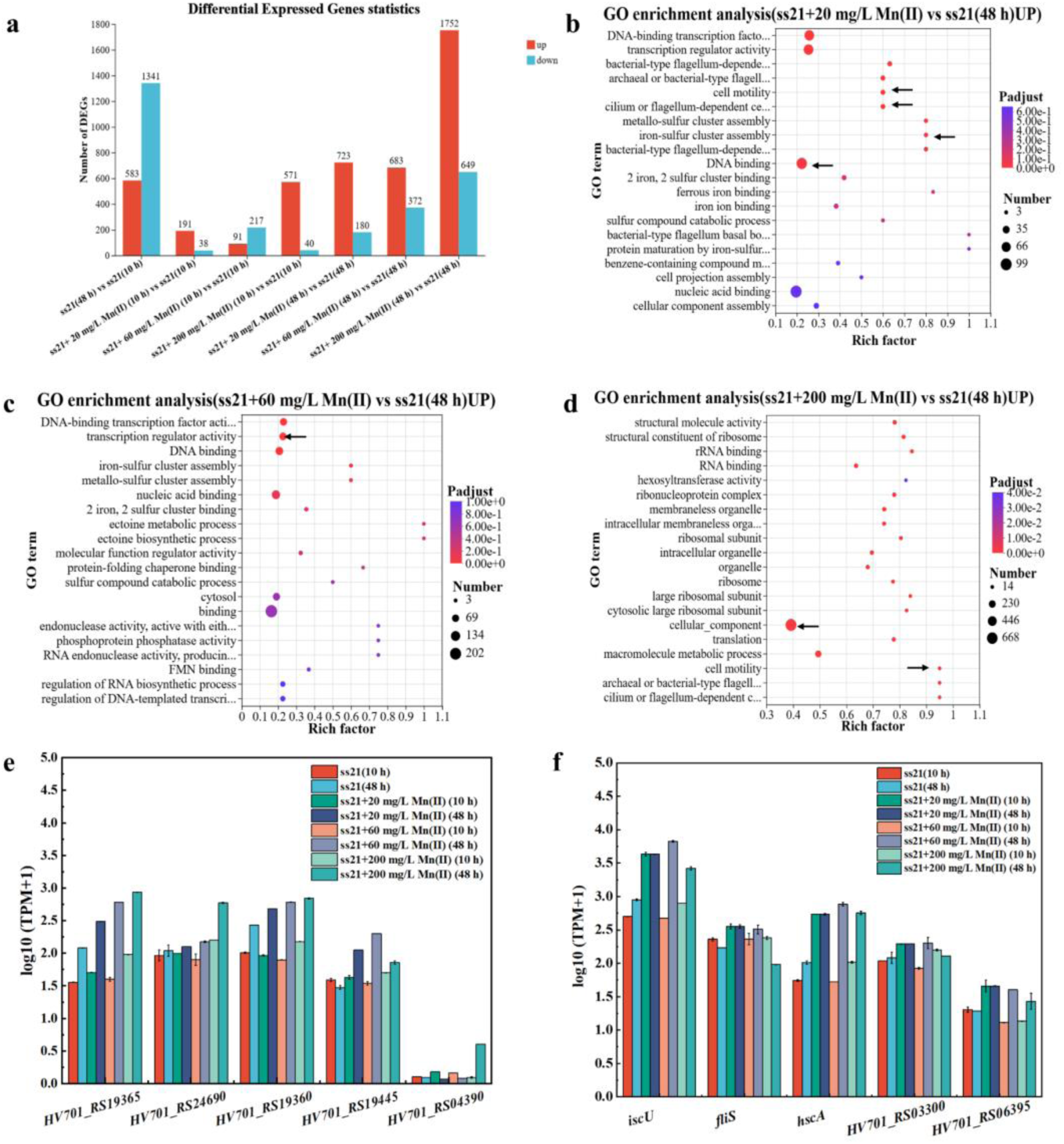
Analysis of differential gene expressions in ss21 with different Mn(Ⅱ) concentration (a), GO functional enrichment enrichment of (b)ss21 + 20 mg/L Mn(Ⅱ) vs ss21 (48 h) UP (b), ss21 + 60 mg/L Mn(Ⅱ) vs ss21 (48 h) UP (c), ss21 + 200 mg/L Mn(Ⅱ) vs ss21 (48 h) UP (d) (The size of bubbles represents the number of genes in the GO enrichment bubble charts); Expressions of genes related to direct (e) and indirect (f) Mn(Ⅱ) oxidation.

To further determine the differentially genes expressed pathways of Mn(Ⅱ) oxidation process, the GO enrichment of the target genes were confirmed. At 48 h, pathway enrichment analysis revealed significant enrichment (FDR<0.05) in pathways under different Mn(Ⅱ) concentration (Fig. 4b, c, and d). For 20 mg/L Mn(Ⅱ), some key pathways enriched, including cilium or flagellum - dependent cell motility (FDR = 0.0023) pathway, cell motility (FDR = 0.0024) pathway, iron-sulfur cluster assembly (FDR = 0.0024) pathway and DNA binding (FDR = 0.0349) pathway (Fig. 4b). Collectively, the enrichment of these pathways suggested that bacterial enhanced motility of cells to increase the contaction with Mn(Ⅱ). Separately, the iron-sulfur cluster assembly pathway contributed to maintain intracellular redox balance under Mn(Ⅱ) stress. In parallel, DNA-binding pathway coordinated the expression of Mn(Ⅱ) oxidation related and stress response genes, thereby synergistically promoting Mn(Ⅱ) oxidation efficiency (33–35). In these enrichment pathways, the genes associated with Mn(Ⅱ) oxidation process were listed in Table S1. Genes involved in cilium or flagellum - dependent cell motility and cell motility pathways, including *flgG*, *flgC*, *flgB*, *fliS, fliE*, and *fliG* were markedly upregulated, these genes effect the flagellar assembly and mobility function of bacteria (36), which promoted bacterial cells to migrate toward Mn(Ⅱ) to accelerate Mn(Ⅱ) oxidation (37). In parallel, the iron-sulfur cluster assembly pathway was enriched, with increased expression of *iscX*, *hscA*, *iscA*, *iscU* and the transcriptional regulator *iscR*. As the iron-sulfur cluster assembly pathway is essential for the generation of redox cofactors controlling the electron transfer and oxidative stress defense, upregulation of these genes suggested the enhanced intracellular redox capacity of the ss21 strain to cope with Mn(Ⅱ) and defense ability for Mn(Ⅱ) stress (38). Moreover, in the DNA-binding pathway, the Cu(I)-responsive transcription factor (*cueR*) associated with the transcriptional expression of multicopper oxidases was upregulated. The upregulation of *cueR* implied the multicopper oxidases were activated to progress Mn(Ⅱ) oxidation by the ss21 strain (39, 40).

For 60 mg/L Mn(Ⅱ), the DNA-binding transcription factor activity (FDR = 0.0361) pathway, and transcription regulator activity (FDR = 0.0361) pathway were significantly upregulated (Fig. 4c). These pathways were accompanied with redox stress regulation, antioxidant defense, and metabolic processes participated in Mn(Ⅱ) oxidation (41, 42). The genes related to Mn(Ⅱ) oxidation in these pathways are listed in Table S2. In the DNA-binding transcription factor activity pathway, *cueR* and *iscR* were significantly expressed which was observed under 20 mg/L Mn(Ⅱ). This result demonstrates that *cueR* and *iscR* are essential for the Mn(Ⅱ) oxidation of the ss21 strain. In the transcription regulator activity pathway, the genes *HV701_RS03300* and *HV701_RS17390* encode MerR family transcriptional regulators that were critical for sensing elevated Mn(Ⅱ) stress and promoting the strain to maintain the physiological homeostasis required for sustained Mn(Ⅱ) oxidation (43). The genes *HV701_RS10460*, *HV701_RS18625*, and *HV701_RS06395*, encoding LysR family transcriptional regulators, which were related to the oxidative stress defense and cellular redox homeostasis (44). The expression of these genes enabled the ss21 strain to effectively oxidze Mn(Ⅱ).

At 200 mg/L Mn(Ⅱ), the significantly upregulated pathways included the cellular component organization (FDR = 0.000934) pathway and cell motility (FDR = 0.000018) pathway (Fig. 4d). Concurrently, the enrichment of the cellular component organization pathway likely supported the repair of cellular structural against the toxicity of Mn(Ⅱ) (45).The genes associated with the Mn(Ⅱ) oxidation process in these pathways are listed in Table S3. In cell motility pathway, *motA*, *flgG*, *fliG*, *fliH*, *fliI* and *fliJ* were markedly upregulated, enhancing the bacterial motility to contact with Mn(Ⅱ) under the stress of high Mn(Ⅱ) concentration (4, 10). Additionally, in the cellular component organization pathway, the gene *HV701_RS19365* encoding an LLM-type flavin oxidoreductase was significantly upregulated, as an electron transfer relay to support extracellular redox process of strain (46). Concurrently, *HV701_RS24690*, encoding a quinone oxidoreductase, drivs the generation of reactive oxygen species (ROS) to facilitate indirect Mn(Ⅱ) oxidation (47, 48). The upregulation of *HV701_RS19360*, the gene encoding the thioredoxin which involved redox enzyme activity (49). Furthermore, the upregulation of *HV701_RS19445* encoding the glutathione peroxidase, indirectly protecting cells from oxidative damage to enhance Mn(Ⅱ) oxidation (50, 51). *HV701_RS04390* gene encodes a copper resistance system multicopper oxidase that likely serves as a key functional enzyme for the direct catalytic oxidation of Mn(Ⅱ) into BioMnOx (52).

The key genes associated with the Mn(Ⅱ) by ss21 strain were obtained from the above GO enrichment analysis (Fig. 4e and f). The genes associated with direct Mn(Ⅱ) oxidation were identified, including *HV701_RS19365*, *HV701_RS24690*, *HV701_RS19360*, *HV701_RS19445* and *HV701_RS04390* (Fig. 4e). Compared with ss21 (48 h), the expression levels of these genes by 1.19, 1.03, 1.10, 1.39, and 0.74 in ss21 + 20 mg/L Mn(Ⅱ) (48 h) under different Mn(Ⅱ) concentration, respectively, and the expression levels of these genes were upregulated by 1.34, 1.07, 1.14, 1.56 and 0.85 folds in ss21 + 60 mg/L Mn(Ⅱ) (48 h), respectively. While in ss21 + 200 mg/L Mn(Ⅱ) (48h), the upregulation increased to 1.41, 1.36, 1.17, 1.26 and 6.27 folds, respectively. Notably, *HV701_RS04390* gene significant upregulated at 200 mg/L Mn(Ⅱ), exhibiting a unique Mn(Ⅱ) concentration response. The expression of these genes contributed to the construction of electron transfer intermediates, extracellular redox processes and production of O₂⁻· to mediate Mn(Ⅱ) oxidation. Moreover, the expression levels of genes associated with indirect Mn(Ⅱ) oxidation, including *iscU*, *fliS*, *hscA*, *HV701_RS03300*, and *HV701_RS06395*, were essentially upregulated (Fig. 4f). Compared with ss21 (48 h), the expression levels of these genes by 1.16, 0.89, 1.37, 1.01 and 1.11 in ss21 + 200 mg/L Mn(Ⅱ) (48 h), respectively. These genes contributed to indirect Mn(Ⅱ) oxidation in strain ss21 by coordinating electron transfer control, oxidative stress defense, bacterial motility, and the maintenance of physiological homeostasis.

KEGG pathways exhibited significant enrichment (p < 0.05) was mapped (Fig. 5). Notably, pathways with Mn(Ⅱ) oxidation process included ribosome, ABC transporters, quorum sensing, biofilm formation-pseudomonas aeruginosa, atrazine degradation, flagellar assembly, and bacterial chemotaxis pathway. Ribosomes pathway are related to the protein synthesis of enzymes for Mn(Ⅱ) oxidation (53, 54). ABC transporters can efficiently import Mn(Ⅱ) from the external environment to provide sufficient Mn(Ⅱ) sources for the bacteria and facilitate the biological Mn(Ⅱ) oxidation process (55). Quorum sensing and biofilm formation-pseudomonas aeruginosa pathways promoted the accumulation of signaling molecules, which are known to be involved in the expression of Mn(Ⅱ) oxidizing genes and biofilm development, thereby enhancing Mn(Ⅱ) tolerance and Mn(Ⅱ) oxidation efficiency (56). Atrazine degradation and biosynthesis of ansamycins pathways has been reported to be involved in the generation of oxidative stress and metabolic inhibition, forcing bacteria to balance the intracellular redox potential through Mn(Ⅱ) transport and Mn(Ⅱ) redox mechanisms, ensuring the normal Mn(Ⅱ) oxidation under oxidative stress and metabolic inhibition (57, 58). Flagellar pathway drived bacterial to move towards Mn(Ⅱ) rich areas, increasing the contaction of bacterial with Mn(Ⅱ) (34, 59).

**Fig. 5.**
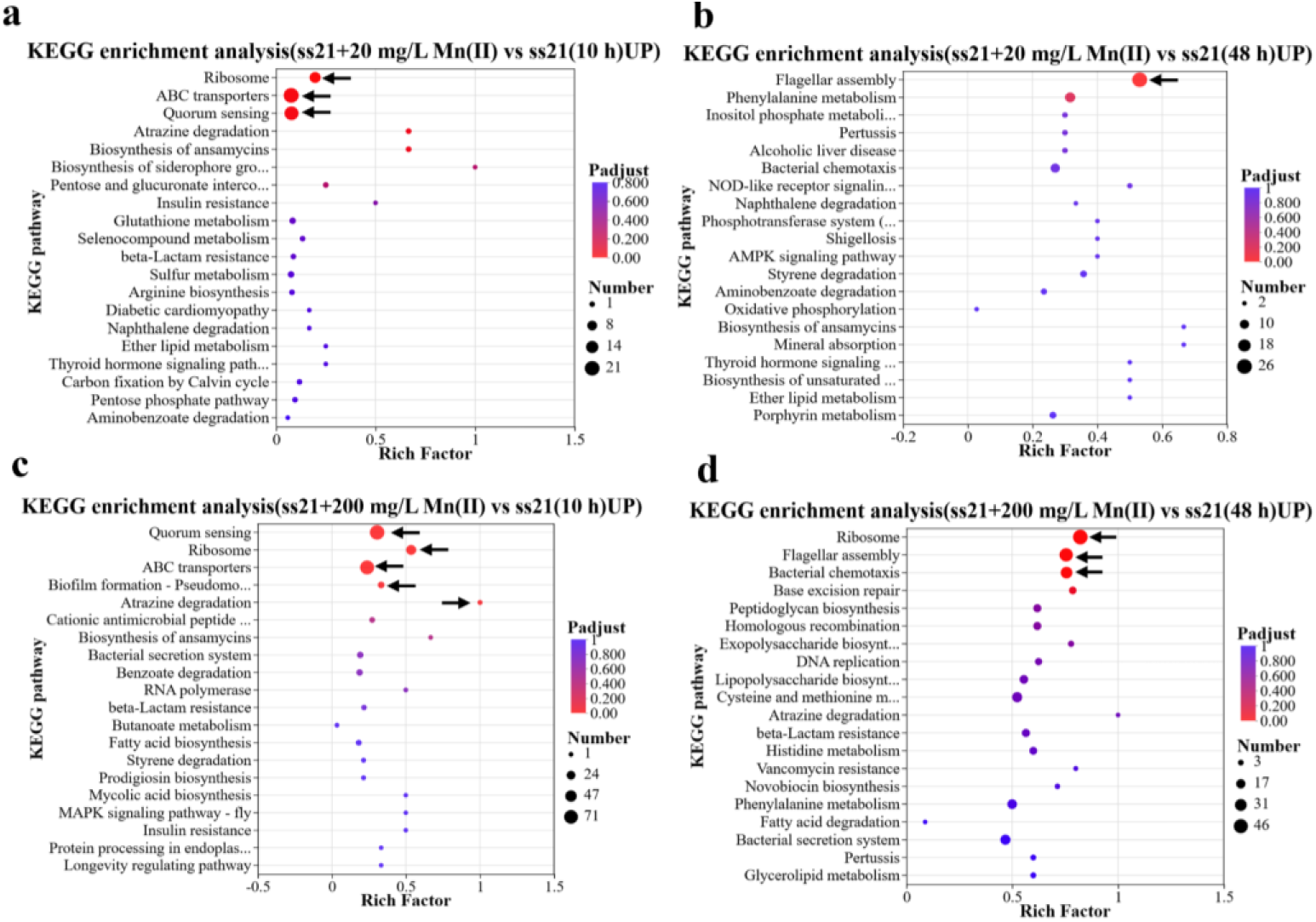
KEGG enrichment analysis of differentially expressed genes in ss21 + 20 mg/L Mn(Ⅱ) vs ss21 (10 h)UP (a) and (48 h)UP (b), ss21 + 60 mg/L Mn(Ⅱ) vs ss21 (10 h)UP (c) and (48 h)UP (d), ss21 + 200 mg/L Mn(Ⅱ) vs ss21 (10 h)UP (e) and (48 h)UP (f).

### METABOLOMICS ANALYSIS

To further investigate the metabolomic variations of the ss21 strain during the Mn(Ⅱ) oxidation process, a non-targeted metabolomic was conducted in the ss21 strain under different initial Mn(Ⅱ) concentrations (Fig. 6). As shown in the PCA analysis (Fig. 6a), all experimental groups were clearly separated with no overlap, indicating that the addition of Mn(Ⅱ) induced distinct alterations in the metabolome of strain ss21. In ss21, 348 metabolites were upregulated and 1020 were downregulated, when comparing 48 h with 10 h (Fig. 6b). At 10 h, strain ss21 exhibited 652, 979, and 1384 differentially expressed metabolites in the ss21+20 mg/L Mn(Ⅱ), ss21+60 mg/L Mn(Ⅱ), and ss21+200 mg/L Mn(Ⅱ) groups, respectively. At 48 h, the strain exhibited 1371, 1229, and 1388 metabolites in the corresponding Mn(Ⅱ) groups, respectively. These results showed that the number of differentially expressed metabolites significantly increased as the initial Mn(Ⅱ) concentration increased from 20 mg/L to 200 mg/L. Furthermore, compared to the 10 h, the number differentially expressed metabolites significantly increased at 48 h. Overall, the higher initial Mn(Ⅱ) concentrations stimulated the expression trend of different metabolites, which consistented with that the srtrain had the highest the Mn(Ⅱ) oxidation efficiency at 200 mg/L Mn(Ⅱ).

**Fig. 6.**
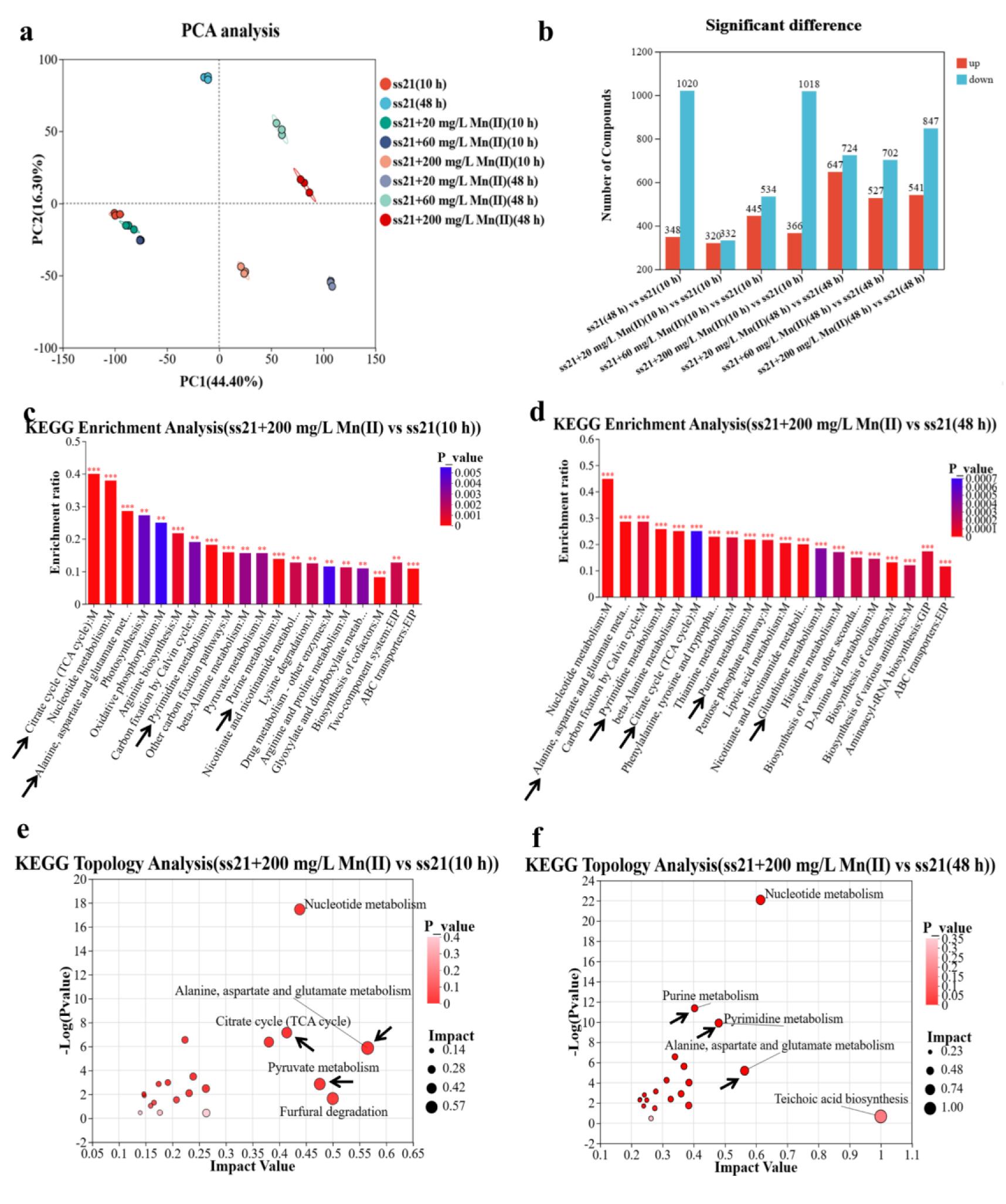
The different metabolites and metabolic pathways in ss21 strain with and without Mn(Ⅱ) (a) PCA analysis; (b) different metabolites. KEGG enrichment analysis of (c) ss21 + 200 mg/L Mn(Ⅱ) vs ss21 (10 h), (d) ss21 + 200 mg/L Mn(Ⅱ) vs ss21 (48 h); KEGG topology analysis of (e) ss21 + 200 mg/L Mn(Ⅱ) vs ss21 (10 h), (f) ss21 + 200 mg/L Mn(Ⅱ) vs ss21(48 h).

KEGG enrichment and topological analysis further revealed the significantly enriched different metabolites (Fig. 6c and d). Enrichment of metabolic pathways are closely related to Mn(Ⅱ) oxidation by ss21 strain, including citrate cycle (TCA cycle), alanine, aspartate, glutamate metabolism, pyrimidine metabolism, and purine metabolism. The enrichment of these pathways also been detected in strain at 20 and 60 mg/L Mn(Ⅱ) (Fig S3 and S4). The TCA cycle has been implicated as a central bioenergetic hub that modulated the production amount of BioMnOx by regulating intracellular carbon flux and energy supply (60). Furthermore, alanine, aspartate, glutamate metabolism play a critical role in pH regulation during the oxidation process, creating an alkaline microenvironment that facilitated the occurrence of Mn(Ⅱ) oxidation (61, 62). The pyrimidine metabolism and purine metabolism play an essential role in supporting bacterial growth during Mn(Ⅱ) oxidation by facilitating nitrogen assimilation from *β*-alanine, and ensuring the biosynthesis of nucleotides and energy carriers (63, 64). Concurrently, These metabolic pathways are critical to electron transfer, Mn(Ⅱ) oxide deposition, pH regulation, and nutrient assimilation during the Mn(Ⅱ) oxidation process.

Cluster heatmap identified the importance and expression trend of different metabolites, and specific metabolites correlated with Mn(Ⅱ) oxidation genes (Fig. 7) “L-Tyrosine” and “L-Isoleucine” regenerated the NADH/NADPH pools required by electron transfer of quinone oxidoreductase, corresponding to the Mn(Ⅱ) oxidation genes *HV701_RS24690* (65, 66). “Glutamic acid” involved in the synthesis of glutathione peroxidase and thioredoxin to scavenge excess ROS generated, and preventing irreversible protein damage from oxidative stress of high Mn(Ⅱ) concentration, corresponding to expression of the Mn(Ⅱ) oxidation genes *HV701_RS19445* and *HV701_RS19360* (67, 68). “Gln-His-His” could alleviate oxidative stress induced by excess Mn(Ⅱ), and guiding Mn(Ⅱ) ions to specific oxidation sites, to enchance Mn(Ⅱ) oxidation process (69). “Flavin Adenine Dinucleotide” (FAD) influenced the activity of LLM-type flavin oxidoreductase that producted by the expression of *HV701_RS19365*, the LLM-type flavin oxidoreductase used FAD as an electronic medium to convert Mn(Ⅱ) to BioMnOx (70–72). “Xanthine” may enhance the activity of *HV701_RS24690*, stimulating the production of ROS. Moreover, xanthine is involved in cellular redox metabolism and promotes the indirect oxidation of Mn(Ⅱ) (73, 74). These representative upregulated metabolites are closely associated with the Mn(Ⅱ) oxidation function of strain ss21 and support its high efficiency Mn(Ⅱ) oxidation.

**Fig. 7.**
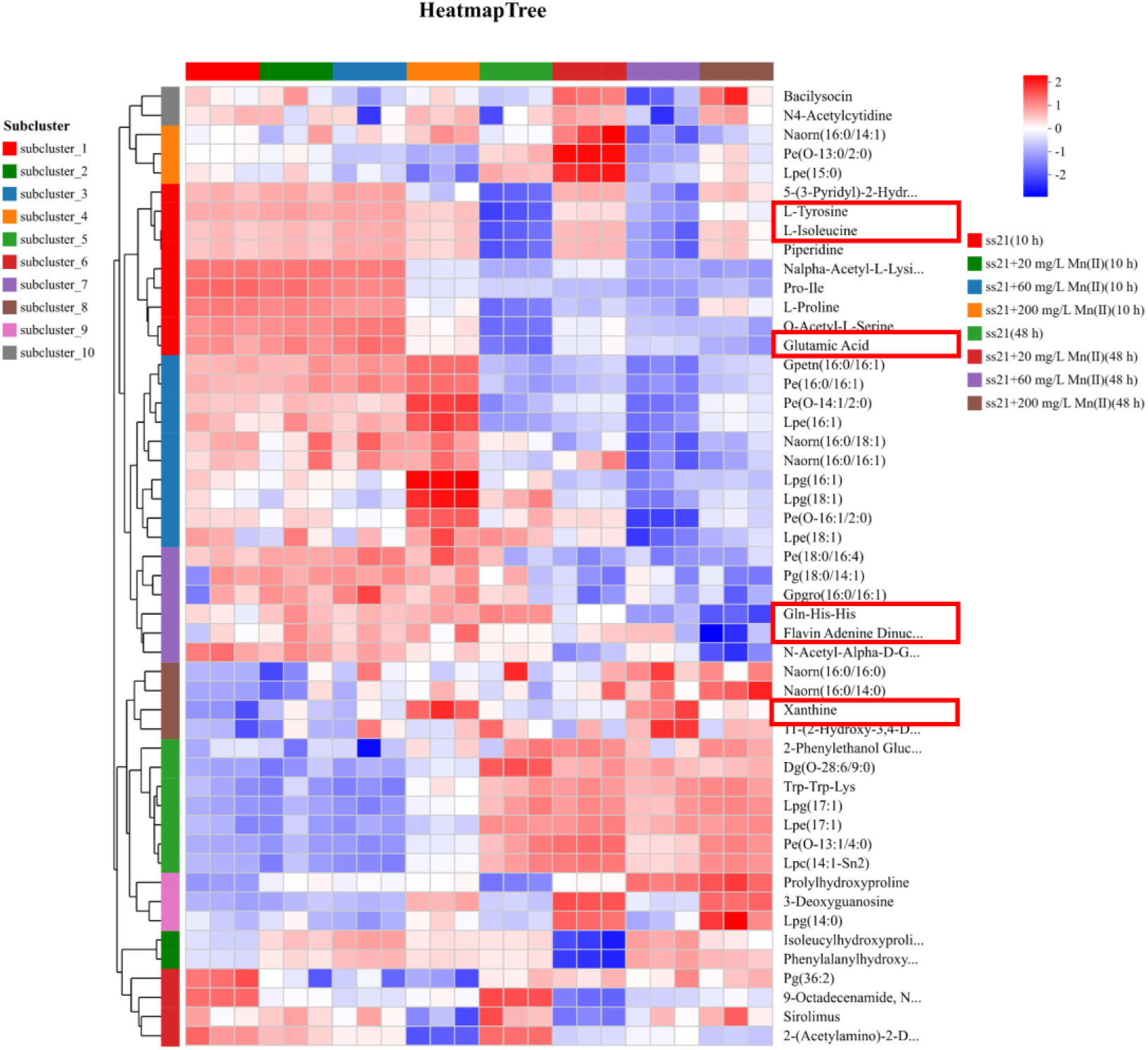
Cluster heatmap of metabolite profiles in ss21 across various Mn(Ⅱ) concentrations.

## CONCLUSION

This study showed that *Achromobacter pulmonis* ss21 achieved a maximum Mn(Ⅱ) oxidation efficiency of 98.82% at an initial Mn(Ⅱ) concentration of 200 mg/L. Integrated transcriptomic and metabolomic analyses revealed that ss21 employs a synergistic mechanism to drive Mn(Ⅱ) oxidation. The *HV701_RS04390*, *HV701_RS19365* and *HV701_RS24690* gene are involved in extracellular electron transfer and ROS generation that supported direct Mn(Ⅱ) oxidation. Concurrently, *HV701_RS19365* and *HV701_RS24690* are closely associated with the enrichment of intracellular metabolites, including L-tyrosine, L-isoleucine, FAD, and xanthine. The ss21 strain upregulated the *HV701_RS19360* and *HV701_RS19445* genes to mitigate the oxidative stress arising from enhanced redox activity to maintain intracellular redox homeostasis, coordinating with glutamic acid and Gln-His-His metabolite. In addition, the *iscU*, *hscA*, *fliS*, *HV701_RS03300*, and *HV701_RS06395* genes controlled oxidative stress defense, bacterial motility, and the maintenance of physiological homeostasis, contributing to indirect Mn(Ⅱ) oxidation. This study expanded the known species of Mn(Ⅱ)-oxidizing bacteria and advanced the understanding of microbial Mn(Ⅱ) oxidation mechanisms in the geochemical element Mn(Ⅱ) cycle.

## DATA AVAILABILITY

All data generated or analyzed during this study are included in this published article and its supplemental material.

---

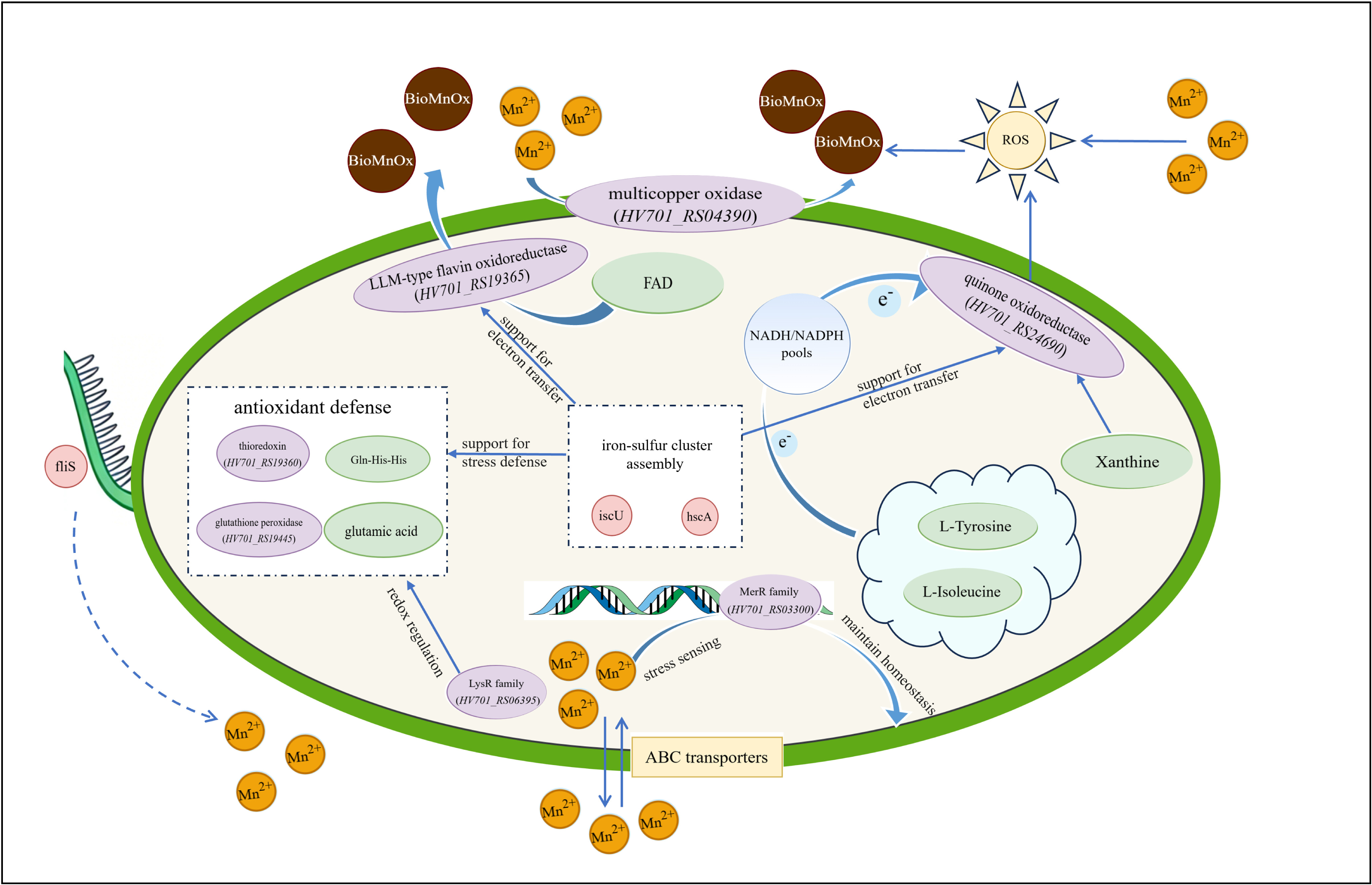

## REFERENCES

1. Hansel CM, Zeiner CA, Santelli CM, Webb SM. 2012. Mn(Ⅱ) oxidation by an ascomycete fungus is linked to superoxide production during asexual reproduction. Proc Natl Acad Sci USA 109:12621–12625. 10.1073/pnas.1203885109.

2. Mo W, Wang H, Wang J, Wang Y, Liu Y, Luo Y, He M, Cheng S, Mei H, He J, Su J. 2024. Advances in Research on Bacterial Oxidation of Mn(Ⅱ): A Visualized Bibliometric Analysis Based on CiteSpace. Microorganisms 12:1611. 10.3390/microorganisms12081611.

3. Bai Y, Su J, Wen Q, Huang T, Chang Q, Ali A. 2021. Characterization and mechanism of Mn(Ⅱ)-based mixotrophic denitrifying bacterium (*Cupriavidus sp.* HY129) in remediation of nitrate (NO_3_^−^-N) and manganese (Mn(Ⅱ)) contaminated groundwater. J Haz Mat 406:124414. 10.1016/j.jhazmat.2020.124414.

4. Yu J, Xu J, Wang H. 2025. Mechanism of Mn(Ⅱ) oxidation by *Bacillus altitudinis* BYD19 under ofloxacin exposure: Insights from transcriptome and metabolomics. J Environ Chem Eng 13:117394. 10.1016/j.jece.2025.117394.

5. Tebo BM, Bargar JR, Clement BG, Dick GJ, Murray KJ, Parker D, Verity R, Webb SM. 2004. BIOGENIC MANGANESE OXIDES: Properties and Mechanisms of Formation. Annu Rev Earth Planet Sci 32:287–328. 10.1146/annurev.earth.32.101802.120213.

6. Li X, Yuan X, Wei Y, He L, Li Y, Qiu M, Liu Y, Dong N, Zhang C, Pang X. 2025. Isolation of a novel manganese-oxidizing bacterium *Lysinibacillus xylanilyticus* M125: characterization, structural evolution, and Cd-adsorption activity of biogenic Mn oxides produced by the strain. Front Microbiol 16:1622784. 10.3389/fmicb.2025.1622784.

7. Zhang Y, Tang Y, Qin Z, Luo P, Ma Z, Tan M, Kang H, Huang Z. 2019. A novel manganese oxidizing bacterium-*Aeromonas hydrophila* strain DS02: Mn(Ⅱ) oxidization and biogenic Mn oxides generation. J Haz Mat 367:539–545. 10.1016/j.jhazmat.2019.01.012.

8. Liu C, Shi B, Guo Y, Wang L, Li S, Zhao C, Zhu L, Wang J, Kim YM, Wang J. 2024. Characteristics of biological manganese oxides produced by manganese-oxidizing bacteria H38 and its removal mechanism of oxytetracycline. Environ Pollut 345:123432. 10.1016/j.envpol.2024.123432.

9. Ou-yahia D, Üstüntürk-Onan M, Ilhan-Sungur E. 2024. Isolation and Identification of a Manganese-Oxidizing Bacterium from Produced Water: Growth and Manganese-Oxidation Ability of *Bacillus zhangzhouensis* in Different Manganese Concentrations. Arab J Sci Eng 49:67–75. 10.1007/s13369-023-08001-6.

10. Huo Y, Mo J, He Y, Twagirayezu G, Xue L. 2022. Transcriptome analysis reveals manganese tolerance mechanisms in a novel native bacterium of *Bacillus altitudinis* strain HM-12. Sci Total Environ 846:157394. 10.1016/j.scitotenv.2022.157394.

11. Andeer PF, Learman DR, McIlvin M, Dunn JA, Hansel CM. 2015. Extracellular haem peroxidases mediate Mn(Ⅱ) oxidation in a marine *Roseobacter* bacterium via superoxide production. Environ Microbiol 17:3925–3936. 10.1111/1462-2920.12893.

12. Geszvain K, McCarthy JK, Tebo BM. 2013. Elimination of Manganese(Ⅱ,Ⅲ) Oxidation in *Pseudomonas putida* GB-1 by a Double Knockout of Two Putative Multicopper Oxidase Genes. Appl Environ Microbiol 79:357–366. 10.1128/AEM.01850-12.

13. Butterfield CN, Soldatova AV, Lee S-W, Spiro TG, Tebo BM. 2013. Mn(Ⅱ,Ⅲ) oxidation and MnO_2_ mineralization by an expressed bacterial multicopper oxidase. Proc Natl Acad Sci USA 110:11731–11735. 10.1073/pnas.1303677110.

14. Soldatova AV, Butterfield C, Oyerinde OF, Tebo BM, Spiro TG. 2012. Multicopper oxidase involvement in both Mn(Ⅱ) and Mn(Ⅲ) oxidation during bacterial formation of MnO_2_. J Biol Inorg Chem 17:1151–1158. 10.1007/s00775-012-0928-6.

15. Francis CA, Tebo BM. 2002. Enzymatic Manganese(Ⅱ) Oxidation by Metabolically Dormant Spores of Diverse *Bacillus* Species. Appl Environ Microbiol 68:874–880. 10.1128/AEM.68.2.874-880.2002.

16. Su J, Bao P, Bai T, Deng L, Wu H, Liu F, He J. 2013. CotA, a Multicopper Oxidase from *Bacillus pumilus* WH4, Exhibits Manganese-Oxidase Activity. PLoS ONE 8:e60573. 10.1371/journal.pone.0060573.

17. El Gheriany IA, Bocioaga D, Hay AG, Ghiorse WC, Shuler ML, Lion LW. 2009. Iron Requirement for Mn(Ⅱ) Oxidation by *Leptothrix discophora* SS-1. Appl Environ Microbiol 75:1229–1235. 10.1128/AEM.02291-08.

18. Ridge JP, Lin M, Larsen EI, Fegan M, McEwan AG, Sly LI. 2007. A multicopper oxidase is essential for manganese oxidation and laccase-like activity in *Pedomicrobium sp.* ACM 3067. Environ Microbiol 9:944–953. 10.1111/j.1462-2920.2006.01216.x.

19. Anderson CR, Johnson HA, Caputo N, Davis RE, Torpey JW, Tebo BM. 2009. Mn(Ⅱ) Oxidation Is Catalyzed by Heme Peroxidases in “*Aurantimonas manganoxydans*” Strain SI85-9A1 and *Erythrobacter sp.* Strain SD-21. Appl Environ Microbiol 75:4130–4138. 10.1128/AEM.02890-08.

20. Gehin G, Carraro N, Gurfield K, Van Der Meer JR, Peña J. 2025. *Pseudomonas putida* Coordinates the Expression of two Manganese Oxidases and Optimizes Manganese Oxide Precipitation in Response to Aqueous Mn(Ⅱ). Environ Sci Technol 59:13777–13786. 10.1021/acs.est.5c00255.

21. Medina M, Rizo A, Dinh D, Chau B, Omidvar M, Juarez A, Ngo J, Johnson HA. 2018. MopA, the Mn Oxidizing Protein From *Erythrobacter sp.* SD-21, Requires Heme and NAD^+^ for Mn(Ⅱ) Oxidation. Front Microbiol 9:2671. 10.3389/fmicb.2018.02671.

22. He Z, Gao J, Li Q, Wei Z, Zhang D, Pan X. 2024. Enhanced oxidation of Mn(Ⅱ) and As(Ⅲ) by aerobic granular sludge via ferrous citrate: Key roles of colloidal iron and extracellular superoxide radical. Water Res 268:122705. 10.1016/j.watres.2024.122705.

23. Yao M-C, Huang Q, Xie H-X, Zhang X, Sheng G-P. 2025. Aquatic Biofilm as a Hotspot for Manganese Oxidation Enabled by Microbial Extracellular Matrix-Mediated Dual-Pathway Electron Transfer. Environ Sci Technol 59:23508–23518. 10.1021/acs.est.5c09864.

24. Li Q, Shi M, Liao Q, Li K, Huang X, Sun Z, Yang W, Si M, Yang Z. 2024. Molecular response to the influences of Cu(Ⅱ) and Fe(Ⅲ) on forming biogenic manganese oxides by *Pseudomonas putida* MnB1. J Haz Mat 477:135298. 10.1016/j.jhazmat.2024.135298.

25. Wang H, Yang Q, Yan Q, Wen Q. 2021. Primary insight into the cathode strengthened electrons transport and nitrous oxide reduction during hydrogenotrophic denitrification in bioelectrochemical system (BES). J Environ Chem Eng 9:104723. 10.1016/j.jece.2020.104723.

26. Li Y, Xu Z, Ma H, Hursthouse AS. 2019. Removal of Manganese(Ⅱ) from Acid Mine Wastewater: A Review of the Challenges and Opportunities with Special Emphasis on Mn-Oxidizing Bacteria and Microalgae. Water 11: 2493. 10.3390/w11122493.

27. Hatayama K. 2020. Manganese Carbonate Precipitation Induced by Calcite-Forming Bacteria. Geomicrobiol J 37:603–609. 10.1080/01490451.2020.1743391.

28. Ma H, Liu X, Wen Z, Yi X, Liu Y, Zhou H. 2023. Competitive Mn(Ⅱ) removal occurs in *Lysinibacillus sp.* MHQ-1 through microbially-induced carbonate precipitation (MICP) and indirect Mn(Ⅱ) oxidation. Environ Res 239:117373. 10.1016/j.envres.2023.117373.

29. Barreto J, Bagus PS, Stavale F. 2024. Multiplet XPS analysis of the Mn 2p for Mn_3_O_4_ thin films. J Phys-Condens Matter 37:045001. 10.1088/1361-648X/ad8b91.

30. Ilton ES, Post JE, Heaney PJ, Ling FT, Kerisit SN. 2016. XPS determination of Mn oxidation states in Mn (hydr)oxides. Appl Surf Sci 366:475–485. 10.1016/j.apsusc.2015.12.159.

31. Klomp R, Żygadłowska OM, Jetten MSM, Oldham VE, van Helmond NAGM, Slomp CP, Lenstra WK. 2025. Dissolved Mn(Ⅲ) is a key redox intermediate in sediments of a seasonally euxinic coastal basin. Biogeosciences 22:751–765. 10.5194/bg-22-751-2025.

32. Tang Y, Webb SM, Estes ER, Hansel CM. 2014. Chromium(Ⅲ) oxidation by biogenic manganese oxides with varying structural ripening. Environ Sci: Process Impacts 16:2127–2136. 10.1039/C4EM00077C.

33. Bosma EF, Rau MH, van Gijtenbeek LA, Siedler S. 2021. Regulation and distinct physiological roles of manganese in bacteria. FEMS Microbiol Rev 45:fuab028. 10.1093/femsre/fuab028.

34. Ran X, Zhu Z, Long H, Tian Q, You L, Wu X, Liu Q, Huang S, Li S, Niu X, Wang J. 2021. Manganese Stress Adaptation Mechanisms of *Bacillus safensis* Strain ST7 From Mine Soil. Front Microbiol 12:758889. 10.3389/fmicb.2021.758889.

35. Ding H. 2025. Iron-sulfur cluster biogenesis and regulation of intracellular iron homeostasis in *Escherichia coli*. Metallomics 17:mfaf040. 10.1093/mtomcs/mfaf040.

36. Wang Y, Xiao L, Ji J, Hassani D, Wu Y, Niu H, Dong Q. 2025. Transcriptomic analysis of mature biofilm and planktonic cells of *Listeria monocytogenes* under nutritional stress. Food Microbiol 132:104859. 10.1016/j.fm.2025.104859.

37. Colin R, Ni B, Laganenka L, Sourjik V. 2021. Multiple functions of flagellar motility and chemotaxis in bacterial physiology. FEMS Microbiol Rev 45:fuab038. 10.1093/femsre/fuab038.

38. Vallières C, Benoit O, Guittet O, Huang M-E, Lepoivre M, Golinelli-Cohen M-P, Vernis L. 2024. Iron-sulfur protein odyssey: exploring their cluster functional versatility and challenging identification. Metallomics 16:mfae025. 10.1093/mtomcs/mfae025.

39. Outten FW, Outten CE, Hale J, O’Halloran TV. 2000. Transcriptional Activation of an *Escherichia coli* Copper Efflux Regulon by the Chromosomal MerR Homologue, CueR. J Biol 275:31024–31029. 10.1074/jbc.M006508200.

40. Su J, Deng L, Huang L, Guo S, Liu F, He J. 2014. Catalytic oxidation of manganese(Ⅱ) by multicopper oxidase CueO and characterization of the biogenic Mn oxide. Water Res 56:304–313. 10.1016/j.watres.2014.03.013.

41. Casamassimi A, Ciccodicola A. 2019. Transcriptional Regulation: Molecules, Involved Mechanisms, and Misregulation. Int J Mol Sci 20:1281. 10.3390/ijms20061281.

42. Hsu P-C, Lu T-C, Hung P-H, Leu J-Y. 2024. Protein moonlighting by a target gene dominates phenotypic divergence of the Sef1 transcriptional regulatory network in yeasts. Nucleic Acids Res 52:13914–13930. 10.1093/nar/gkae1147.

43. Fang C, Zhang Y. 2022. Bacterial MerR family transcription regulators: activation by distortion. Acta Biochim Biophys Sin 54:25–36. 10.3724/abbs.2021003.

44. Schmidt JJ, Brandenburg VB, Elders H, Shahzad S, Schäkermann S, Fiedler R, Knoke LR, Pfänder Y, Dietze P, Bille H, Gärtner B, Albin LJ, Leichert LI, Bandow JE, Hofmann E, Narberhaus F. 2025. Two redox-responsive LysR-type transcription factors control the oxidative stress response of Agrobacterium tumefaciens. Nucleic Acids Res 53:gkaf267. 10.1093/nar/gkaf267.

45. Ciancio Casalini L, Piazza A, Masotti F, Garavaglia BS, Ottado J, Gottig N. 2022. Manganese oxidation counteracts the deleterious effect of low temperatures on biofilm formation in *Pseudomonas sp.* MOB-449. Front Mol Biosci 9. 10.3389/fmolb.2022.1015582.

46. Light SH, Su L, Rivera-Lugo R, Cornejo JA, Louie A, Iavarone AT, Ajo-Franklin CM, Portnoy DA. 2018. A flavin-based extracellular electron transfer mechanism in diverse Gram-positive bacteria. Nature 562:140–144. 10.1038/s41586-018-0498-z.

47. Abou-Hamdan A, Mahler R, Grossenbacher P, Biner O, Sjöstrand D, Lochner M, Högbom M, von Ballmoos C. 2022. Functional design of bacterial superoxide:quinone oxidoreductase. Bba-bioenergetics 1863:148583. 10.1016/j.bbabio.2022.148583.

48. Learman DR, Voelker BM, Vazquez-Rodriguez AI, Hansel CM. 2011. Formation of manganese oxides by bacterially generated superoxide. Nat Geosci 4:95–98. 10.1038/ngeo1055.

49. Lee S, Kim SM, Lee RT. 2013. Thioredoxin and Thioredoxin Target Proteins: From Molecular Mechanisms to Functional Significance. Antioxid Redox Signal 18:1165–1207. 10.1089/ars.2011.4322.

50. Carmel-Harel O, Storz G. 2000. Roles of the Glutathione- and Thioredoxin-Dependent Reduction Systems in the *Escherichia Coli* and *Saccharomyces Cerevisiae* Responses to Oxidative Stress. Annu Rev Microbiol 54:439–461. 10.1146/annurev.micro.54.1.439.

51. Lu J, Holmgren A. 2014. The thioredoxin antioxidant system. Free Radical Biol Med 66:75–87. 10.1016/j.freeradbiomed.2013.07.036.

52. Butterfield CN, Tao L, Chacón KN, Spiro TG, Blackburn NJ, Casey WH, Britt RD, Tebo BM. 2015. Multicopper manganese oxidase accessory proteins bind Cu and heme. Bba-proteins Proteom 1854:1853–1859. 10.1016/j.bbapap.2015.08.012.

53. Cui Z, Li X, Shin J, Gamper H, Hou Y-M, Sacchettini JC, Zhang J. 2022. Interplay between an ATP-binding cassette F protein and the ribosome from *Mycobacterium tuberculosis*. Nat Commun 13:432. 10.1038/s41467-022-28078-1.

54. Ousalem F, Singh S, Chesneau O, Hunt JF, Boël G. 2019. ABC-F proteins in mRNA translation and antibiotic resistance. Res Microbiol 170:435–447. 10.1016/j.resmic.2019.09.005.

55. Green RT, Todd JD, Johnston AWB. 2013. Manganese uptake in marine bacteria; the novel MntX transporter is widespread in Roseobacters, Vibrios, Alteromonadales and the SAR11 and SAR116 clades. ISME J 7:581–591. 10.1038/ismej.2012.140.

56. Huang Z, Wang X, Yang L, Palomo A, Liu B, Qin C, Yang J, Andrews CB, Zheng C. 2025. Quorum sensing enhances Mn(Ⅱ) oxidation by *Pseudomonas sp.* YoHu13 via elevated Mn(Ⅲ) intermediate formation. J Haz Mat 501:140735. 10.1016/j.jhazmat.2025.140735.

57. Huang Q, Zhang X, Guo Z, Fu X, Zhao Y, Kang Q, Bai L. 2023. Biosynthesis of ansamitocin P-3 incurs stress on the producing strain *Actinosynnema pretiosum* at multiple targets. Commun Biol 6:860. 10.1038/s42003-023-05227-w.

58. Zhang Y, Meng D, Wang Z, Guo H, Wang Y, Wang X, Dong X. 2012. Oxidative stress response in atrazine-degrading bacteria exposed to atrazine. J Haz Mat 229-230:434–438. 10.1016/j.jhazmat.2012.05.054.

59. Wang Z, Wang J, Liu J, Chen H, Li M, Li L. 2017. Mechanistic insights into manganese oxidation of a soil-borne Mn(Ⅱ)-oxidizing *Escherichia coli* strain by global proteomic and genetic analyses. Sci Rep 7:1352. 10.1038/s41598-017-01552-3.

60. Gu T, Tong Z, Zhang X, Wang Z, Zhang Z, Hwang T-S, Li L. 2022. Carbon Metabolism of a Soilborne Mn(Ⅱ)-Oxidizing *Escherichia coli* Isolate Implicated as a Pronounced Modulator of Bacterial Mn Oxidation. Int J Mol Sci 23:5951. 10.3390/ijms23115951.

61. Feehily C, Karatzas KAG. 2013. Role of glutamate metabolism in bacterial responses towards acid and other stresses. J Appl Microbiol 114:11–24. 10.1111/j.1365-2672.2012.05434.x.

62. Kastenmüller G, Schenk ME, Gasteiger J, Mewes H-W. 2009. Uncovering metabolic pathways relevant to phenotypic traits of microbial genomes. Genome Biol 10:R28. 10.1186/gb-2009-10-3-r28.

63. Yin J, Wei Y, Liu D, Hu Y, Lu Q, Ang EL, Zhao H, Zhang Y. 2019. An extended bacterial reductive pyrimidine degradation pathway that enables nitrogen release from β-alanine. J Biol Chem 294:15662–15671. 10.1074/jbc.RA119.010406.

64. Yoo HC, Yu YC, Sung Y, Han JM. 2020. Glutamine reliance in cell metabolism. Exp Mol Med 52:1496–1516. 10.1038/s12276-020-00504-8.

65. Li X, Zhang H. 2024. Amino acid metabolism, redox balance and epigenetic regulation in cancer. FEBS J 291:412–429. 10.1111/febs.16803.

66. Lu D, Grant M, Lim BL. 2025. NAD(H) and NADP(H) in plants and mammals. Mol Plant 18:938–959. 10.1016/j.molp.2025.05.004.

67. Orlowska K, Fling RR, Nault R, Schilmiller AL, Zacharewski TR. 2023. Cystine/Glutamate Xc^−^ Antiporter Induction Compensates for Transsulfuration Pathway Repression by 2,3,7,8-Tetrachlorodibenzo-*p*-dioxin (TCDD) to Ensure Cysteine for Hepatic Glutathione Biosynthesis. Chem Res Toxicol 36:900–915. 10.1021/acs.chemrestox.3c00017.

68. Vašková J, Kočan L, Vaško L, Perjési P. 2023. Glutathione-Related Enzymes and Proteins: A Review. Molecules 28:1447. 10.3390/molecules28031447.

69. Shalev DE. 2022. Studying Peptide-Metal Ion Complex Structures by Solution-State NMR. IJMS 23:15957. 10.3390/ijms232415957.

70. Zeiner CA, Purvine SO, Zink E, Wu S, Paša-Tolić L, Chaput DL, Santelli CM, Hansel CM. 2021. Mechanisms of Manganese(Ⅱ) Oxidation by Filamentous Ascomycete Fungi Vary With Species and Time as a Function of Secretome Composition. Front Microbiol 12:610497. 10.3389/fmicb.2021.610497.

71. Deng Y, Zhou Q, Wu Y, Chen X, Zhong F. 2022. Properties and Mechanisms of Flavin-Dependent Monooxygenases and Their Applications in Natural Product Synthesis. Int J Mol Sci 23:2622. 10.3390/ijms23052622.

72. Guan Y, Chen X. 2023. Recent Applications of Flavin-Dependent Monooxygenases in Biosynthesis, Pharmaceutical Development, and Environmental Science. Catalysts 13:1495. 10.3390/catal13121495.

73. Battelli MG, Polito L, Bortolotti M, Bolognesi A. 2016. Xanthine Oxidoreductase-Derived Reactive Species: Physiological and Pathological Effects. Oxid Med Cell Longev 2016:3527579. 10.1155/2016/3527579.

74. Jofré I, Matus F, Mendoza D, Nájera F, Merino C. 2021. Manganese-Oxidizing Antarctic Bacteria (Mn-Oxb) Release Reactive Oxygen Species (ROS) as Secondary Mn(Ⅱ) Oxidation Mechanisms to Avoid Toxicity. Biology 10:1004. 10.3390/biology10101004.

